# Neural substrates of verbal repetition deficits in primary progressive aphasia

**DOI:** 10.1101/2020.10.27.357038

**Authors:** Hilary E. Miller, Claire Cordella, Jessica A. Collins, Rania Ezzo, Megan Quimby, Daisy Hochberg, Jason A. Tourville, Bradford C. Dickerson, Frank H. Guenther

**Affiliations:** Department of Speech, Language, & Hearing Sciences, Boston University, Boston, MA 02215; Frontotemporal Disorders Unit, Department of Neurology, Massachusetts General Hospital & Harvard Medical School, Charlestown, MA 02129; Martinos Center for Biomedical Imaging, Department of Radiology, Massachusetts General Hospital, Charlestown, MA 02129; Department of Biomedical Engineering, Boston University, Boston, MA 02215; The Picower Institute for Learning and Memory, Massachusetts Institute of Technology, Cambridge, MA 02139

**Keywords:** primary progressive aphasia, phonological working memory, GODIVA model, repetition, cortical thickness

## Abstract

In this cross-sectional study, we examined the relationship between cortical thickness and performance on several verbal repetition tasks in a cohort of patients with primary progressive aphasia (PPA) in order to test predictions generated by theoretical accounts of phonological working memory (PWM) that predict phonological content buffers in left posterior inferior frontal sulcus (pIFS) and supramarginal gyrus (SMG). Cortical surfaces were reconstructed from magnetic resonance imaging scans from 42 participants diagnosed with PPA. Cortical thickness was measured in a set of anatomical regions spanning the entire cerebral cortex. Correlation analyses were performed between cortical thickness and average score across three PWM related tasks: the Repetition subtest from the Western Aphasia Battery, a forward digit span task, and a backward digit span task. Significant correlations were found between average PWM score across tasks and cortical thickness in left SMG and left pIFS, in support of prior theoretical accounts of PWM. Exploratory whole-brain correlation analyses performed for each of the three behavioral tasks individually revealed a distinct set of positively-correlated regions for each task. Comparison of cortical thickness measures from different PPA subtypes to cortical thickness in age-matched controls further revealed unique patterns of atrophy in the different PPA subtypes.

## Introduction

Primary progressive aphasia (PPA) is a neurodegenerative syndrome, usually arising from Alzheimer’s disease or Frontotemporal Lobar Degeneration, in which language impairment is the most prominent and initial presenting feature (Mesulam, 2003). Subtypes of PPA further characterize specific patterns of language impairment and expected disease progression (Mesulam *et al*., 2009; Gorno-Tempini *et al*., 2011): *semantic-variant PPA (svPPA)* patients demonstrate anomia and impaired single word comprehension; *nonfluent-variant PPA (nfvPPA)* patients demonstrate agrammatism with or without co-occurring apraxia of speech; and *logopenic-variant PPA (lvPPA)* patients demonstrate deficits in lexical retrieval and phonological processing. Cortical thickness measures reveal differential patterns of atrophy across PPA variants (Mesulam *et al*., 2009, 2012; Rohrer *et al*., 2010; Sapolsky *et al*., 2010; Rogalski *et al*., 2014; Collins *et al*., 2017), and have been employed to identify neural regions underlying core speech and language domains including articulatory rate, fluency, and semantic and syntactic processing (Sapolsky *et al*., 2010; Rogalski *et al*., 2011; Mesulam *et al*., 2015; Cordella *et al*., 2019). For example, the characteristic anterior temporal atrophy in svPPA is associated with single-word comprehension abilities, while distinctive left inferior frontal atrophy in nfvPPA correlates with measures of syntactic processing (Amici *et al*., 2007; Sapolsky *et al*., 2010; Rogalski *et al*., 2011). LvPPA is associated with cortical thinning in the temporoparietal junction, with atrophy here also correlated with sentence repetition abilities (Amici *et al*., 2007; Rogalski *et al*., 2011; Lukic *et al*., 2019).

Verbal repetition tasks, such as sentence repetition and digit span tasks, are often used for clinical characterization of the core phonological impairment in lvPPA (Gorno-Tempini *et al*., 2008, 2011; Foxe *et al*., 2013; Meyer *et al*., 2015). However, contradictory evidence suggests that these tasks may not always differentiate lvPPA from other PPA variants or Alzheimer’s disease (Leyton *et al*., 2014; Beales *et al*., 2019). Differences in the various verbal repetition tasks used across studies likely contribute to the divergent results. Thus, further investigation of the neural bases of phonological working memory is critical to differentiate underlying neural mechanisms predictive of repetition impairment in PPA patients on common diagnostic tasks.

In this study, we focus on working memory-related predictions based on the Gradient Order Directions into Velocities of Articulators (GODIVA) model, which is a neurocomputational model of the processes involved in the planning and sequencing of multisyllabic utterances (Bohland *et al*., 2010; Guenther, 2016). According to the model, the content of an upcoming utterance is temporarily stored in two distinct sub-regions of prefrontal cortex: a *metrical structure buffer* in bilateral pre-supplementary motor area (preSMA) and a *phonological content buffer* in left posterior inferior frontal sulcus (pIFS; Bohland & Guenther, 2006). The phonological content buffer is responsible for buffering of phonemes in working memory while earlier portions of the utterance are being articulated. We further posit that the phonological content buffer in left pIFS is distinct from a second phonological buffer located in the left supramarginal gyrus (SMG) that is heavily involved in speech perception and language recognition, as shown in Figure 1.

**Figure 1.**
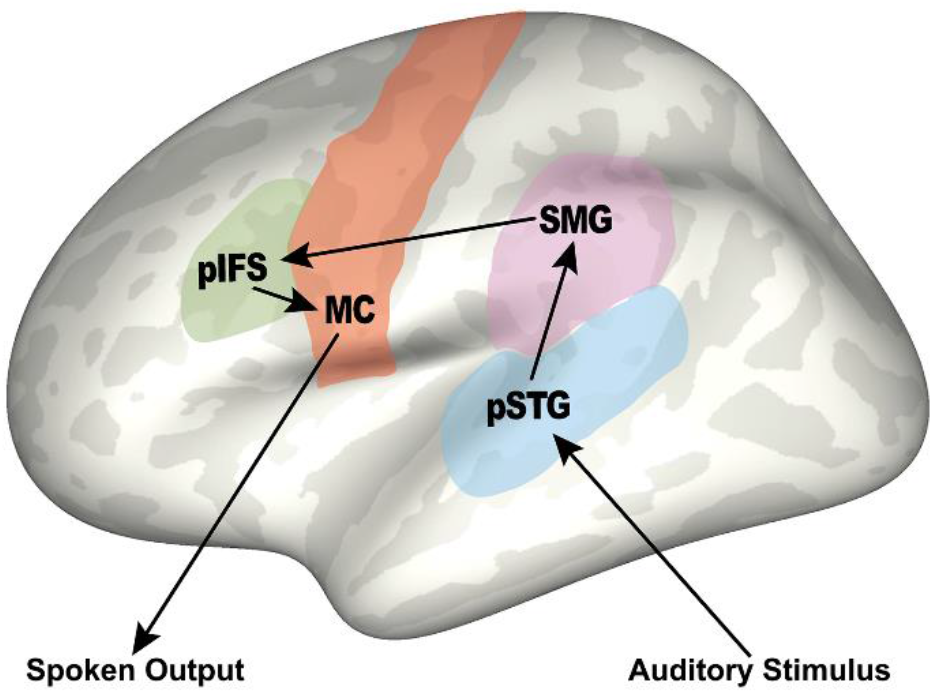
A simplified account of neural processing in verbal repetition tasks showing key left hemisphere brain regions involved: auditory perception in posterior superior temporal gyrus (pSTG), phonological content buffers in supramarginal gyrus (SMG) and posterior inferior frontal sulcus (pIFS), and generation of movement commands in motor cortex (MC), resulting in spoken output of the presented auditory stimulus.

Previous studies investigating repetition in PPA support the involvement of this temporoparietal phonological buffer in verbal repetition tasks (Amici *et al*., 2007; Rogalski *et al*., 2011; Leyton *et al*., 2012; Lukic *et al*., 2019). The current study seeks to extend this work to test for the involvement of an additional phonological content buffer in left pIFS in sentence repetition and digit span working memory tasks, as predicted by GODIVA model. Successful repetition of sentences or digit sequences during these tasks requires accurate buffering and sequencing of each phoneme for subvocal rehearsal and eventual spoken output. The proposed phonological content buffer in left pIFS should therefore be heavily involved in these diagnostic tasks. We also include exploratory whole-brain analyses for each of the three repetition tasks to compare the neural correlates of each task.

## Methods

The study was approved by the Partners Human Research Committee, the Institutional Review Board of Partners HealthCare. All participants provided written informed consent prior to enrollment in the study.

### Diagnostic criteria

Participants included forty-two patients with a diagnosis of PPA selected from the PPA Longitudinal Cohort of the Massachusetts General Hospital (MGH) Frontotemporal Disorders Unit’s Primary Progressive Aphasia Program. For the purposes of the current study, PPA participant selection criteria were (1) an assessment of repetition and working memory (digit span) performance, (2) the availability of an MRI scan, and (3) right-handedness. Fifty-one patients from the PPA Longitudinal Cohort were considered for eligibility, with seven patients excluded due to left-handedness and two due to low quality imaging data. Power calculations indicated that our sample size was adequate to detect a medium strength brain-behavior correlation (*r* = 0.40) similar to those reported previously in PPA (e.g. Cordella *et al*., 2019; Petroi *et al*., 2020), assuming a power of 0.80 and alpha level of 0.05 (one-tailed).

Participants in this cohort undergo a comprehensive clinical evaluation as previously described (Sapolsky *et al*., 2011, 2014), with diagnosis of PPA and subsequent subtype classification made by consensus by the neurologist in consultation with the speech-language pathologist. For each participant, we perform an extensive multidisciplinary assessment including a structured interview of the participant by a neurologist or psychiatrist covering cognition, mood/behavior, sensorimotor function, and daily activities; a neurologic examination, including office-based cognitive testing (for cases in this report, BCD); a speech-language assessment performed by a speech-language pathologist (for cases in this report, MQ or DH), including the Progressive Aphasia Severity Scale (PASS) to specifically assess language impairment from a patient’s premorbid baseline (Sapolsky *et al*., 2014); an MRI scan with T1- and T2-weighted sequences inspected visually by a neurologist. For each participant, a clinician also performs a structured interview with an informant who knows the participant well, augmented with standard questionnaires. For most of the participants in this report, the protocol included the National Alzheimer’s Coordinating Center (NACC) Uniform Data Set measures (using version 2.0 previously and currently version 3.0), as well as supplementary measures. Cases selected for this study had been diagnosed with PPA according to consensus guidelines (Mesulam, 2001; Gorno-Tempini *et al*., 2011). In accordance with these criteria, all participants exhibited a progressive language impairment with a relative preservation of other cognitive functions. Visual inspection of a clinical MRI ruled out other causes of focal brain damage. No participants harbored a pre-existing psychiatric disorder, other neurological disorder or developmental cognitive disorder. This study included nonfluent-variant PPA patients (nfvPPA; N=13), logopenic-variant PPA patients (lvPPA; N=14), and semantic-variant PPA patients (svPPA; N=15). For 10 of the 13 patients diagnosed with nfvPPA, both of the two primary inclusion criteria (i.e., apraxia of speech and agrammatism) were met (with two presenting only with agrammatism and one with only apraxia of speech). Detailed speech/language characteristics per diagnostic group are reported in Table 1.

**Table 1.** Demographic and speech/language characteristics by group, for logopenic variant (lvPPA), nonfluent variant (nfvPPA), and semantic variant (svPPA) patients. CDR = Clinical Dementia Rating; PASS = Progressive Aphasia Severity Score; SD = standard deviation. Differences in group means (post-hoc t-test, p < 0.05) from ^a^lvPPA, ^b^nfvPPA, ^c^svPPA ^d^Indicates time in years between diagnosis date and initial study visit

|  | **<u>lvPPA</u>**<br>**<u>(n = 14)</u>** | **<u>nfvPPA</u>**<br>**<u>(n = 13)</u>** | **<u>svPPA</u>**<br>**<u>(n = 15)</u>** |
| --- | --- | --- | --- |
| **Female, number (%)** | 8 F (57%) | 6 F (46%) | 9 F (60%) |
| **Age, y (SD)** | 71.3 (8.1) | 69.4 (8.4) | 64.7 (7.3) |
| **Education, y (SD)** | 16.2 (3.2) | 15.8 (3.4) | 16.3 (1.9) |
| **Time from Diagnosis<sup>d</sup>, y (SD)** | 0.7 (1.1) | 1.0 (2.3) | 0.9 (1.1) |
| **Mean CDR Language Box Score<sup>e</sup> (SD)** | 0.8 (0.4) | 0.8 (0.6) | 1.1 (0.5) |
| **Mean PASS subdomain scores<sup>e</sup> (SD)** |  |  |  |
| **Articulation** | 0.1 (0.3)<sup>b</sup> | 1.0 (0.9)<sup>a,c</sup> | 0.0 (0.1)<sup>b</sup> |
| **Fluency** | 0.5 (0.3)<sup>b</sup> | 1.0 (0.6)<sup>a,c</sup> | 0.2 (0.3)<sup>b</sup> |
| **Syntax** | 0.4 (0.3) | 0.7 (0.5)<sup>c</sup> | 0.3 (0.3)<sup>b</sup> |
| **Word Retrieval** | 1.0 (0.4)<sup>b</sup> | 0.6 (0.2)<sup>a,c</sup> | 1.1 (0.5)<sup>b</sup> |
| **Repetition** | 0.8 (0.3)<sup>b,c</sup> | 0.4 (0.3)<sup>a</sup> | 0.3 (0.3)<sup>a</sup> |
| **Auditory Comprehension** | 0.7 (0.5) | 0.4 (0.3) | 0.5 (0.5) |
| **Single-Word Comprehension** | 0.2 (0.2)<sup>c</sup> | 0.0 (0.1)<sup>c</sup> | 1.0 (0.5)<sup>a,b</sup> |
| **Mean PASS score, combined subtests** | 0.5 (0.2) | 0.6 (0.3) | 0.5 (0.2) |
| **WAB- Repetition score (SD)** | 71.3 (15.2)<sup>b,c</sup> | 88.8 (14.0)<sup>a</sup> | 86.5 (7.6)<sup>c</sup> |
| **Forward Digit Span score (SD)** | 4.2 (1.6)<sup>c</sup> | 5.5 (1.3) | 6.7 (1.2)<sup>a</sup> |
| **Backward Digit Span score (SD)** | 2.5 (1.6)<sup>c</sup> | 3.3 (0.9) | 4.0 (1.6)<sup>a</sup> |

### Behavioral measures

Participants completed the Repetition subtest of the Western Aphasia Battery-Revised (WAB-R; Kertesz, 2007), which included 15 stimuli items that vary in length. The subtest includes seven single words (1-3 syllables), and eight phrases/sentences (5-14 syllables). Each word, phrase or sentence was read aloud to the participant and participants were instructed to repeat. Points per item were determined based on the standardized scoring guidelines. Points were deducted for omissions of phonemes, syllables or entire words, as well as for phonemic substitutions and additions. Points were not deducted in the case of a timely self-correction of phonemic error or an intelligible sound distortion (i.e., motor speech impairment was not penalized). Stimuli were eligible for scoring only after the first administration. In the current study, the reported total score for the WAB-Repetition subtest refers to the overall percent correct (out of 100 possible points) across all stimuli.

Participants also completed Digit Span Forward and Digit Span Backward subtests from the Uniform Data Set (UDS v3.0) neuropsychological test battery (Weintraub *et al*., 2009). The Digit Span subtests each comprise 14 stimuli digit sets varying in span length (3-9 digits for

Forward subtest; 2-8 digits for Backward subtest). For each span length, there are two stimuli digit sets. Each digit set was read aloud to the participant and the participant was instructed to repeat those numbers in either the exact order they heard them (Digit Span Forward) or to repeat them back in the reverse order (Digit Span Backward). Responses for each digit set were scored as correct/incorrect, and no partial points were awarded. In these subtests, patients were not penalized for either phonological or articulatory errors, provided that the response was intelligible. Testing was discontinued after two consecutive failures on the same span length. In this study, the reported total score refers to the length of the longest correctly repeated sequence. If a participant was unable to correctly repeat the shortest length sequence at least once, they received a total score of zero.

### Structural MRI acquisition and analysis

Imaging data for all PPA patients were acquired on a 3T Magnetom Tim Trio system at MGH (Siemens), using a 12-channel phased-array head coil. For each patient a structural image was obtained using a standard T1-weighted 3D MPRAGE sequence that varied slightly across individuals. 19 patients were scanned using the following parameters: TR = 2530.00 ms, TE = 3.48 ms, flip angle = 7.00 degrees, number of interleaved sagittal slices = 176, matrix dimensions = 256 × 256, FOV = 256 mm, voxel size = 1.00 mm isotropic. Nine patients had the parameters: TR = 2530.00 ms, TE = 1.64 ms, flip angle = 7.00 degrees, number of interleaved sagittal slices = 176, matrix dimensions = 256 × 256, FOV = 256 mm, voxel size = 1.00 mm isotropic. Five patients were scanned with TR = 2300.00 ms, TE = 2.98 ms, flip angle = 9.00 degrees, number of interleaves sagittal slices = 160, matrix = 240 × 256, FOV = 256 mm, voxel size = 1.00 mm isotropic; and two patients had identical parameters with the exception of number of interleaved sagittal slices = 192. Two patients were scanned with TR = 2530.00 ms, TE = 1.61, flip angle = 7.00 degrees, number of interleaved sagittal slices = 208, matrix dimensions = 256 × 256, FOV = 256, mm, voxel size = 1.00 mm isotropic; and one patient had identical parameters with the exception matrix = 280 × 280 FOV = 280 mm, and TE = 1.63 ms. Three patients were scanned with TR = 2200.00 ms, TE = 1.54 ms, flip angle = 7.00 degrees, number of interleaved sagittal slices = 144, matrix = 192 x 192, FOV = 230 mm, voxel size = 1.198mm x 1.198mm x 1.200mm. One remaining patient was scanned with the following: TR = 2400.00 ms, TE = 2.22 ms, flip angle = 8.00 degrees, number of interleaved sagittal slices = 208, matrix dimensions = 300 × 320, FOV = 256 mm, voxel size = 0.80 mm isotropic.

MRI structural images were also obtained for age-matched control participants who did not exhibit any cognitive impairment (n = 25; mean age = 67.4 years, SD = 4.9; 12 female). MRI data for control participants were obtained using the following scan parameters: TR = 2300.00 ms, TE = 2.95 ms, flip angle = 9.00 degrees, number of interleaved sagittal slices = 176, matrix dimensions = 256 x 256, FOV = 270 mm, voxel size = 1.1mm x 1.1mm x 1.200mm.

Cortical reconstructions were generated for each participant’s T1-weighted image using FreeSurfer version 6.0 (https://surfer.nmr.mgh.harvard.edu, Dale, Fischl, & Sereno, 1999; Fischl & Dale, 2000; Fischl, Sereno, & Dale, 1999; Salat et al., 2004). This method has been shown to be reliable in older adults for both spatial localization and absolute magnitude of measurements across multiple scan sessions for the identification of brain-behavior relationships (Dickerson *et al*., 2008). Each cortical reconstruction was inspected for accuracy and any errors in the grey/white matter boundary or pial surface segmentation were manually corrected. Each patient’s reconstructed cortical surface was then parcellated using the SpeechLabel cortical labeling system, which parcellates each hemisphere into 66 anatomically-based regions-of-interest (ROIs) for fine-scale subdivision of cortical regions involved in the speech network and is described in greater detail in previous work from our labs (Cai *et al*., 2014; Cordella *et al*., 2019). Average cortical thickness within each ROI of the SpeechLabel atlas was calculated for each patient.

To identify ROIs demonstrating significant atrophy for each PPA variant compared to controls, independent-sample one-tailed t-tests were conducted for each ROI, using a one-tailed statistical threshold of *p* < 0.001 with FDR corrections. Separate ANOVA analyses were completed to identify differences in cortical thickness in the hypothesized phonologic buffer ROIs in left pIFS and left posterior supramarginal gyrus (pSMG) between each PPA variant.

### Experimental design and statistical analysis

A principal components analysis was first performed for the three working memory scores, which revealed that the three working memory measures contributed essentially equally to the first principal component (coeff = 0.56, 0.60, 0.57 respectively for backward digit span, forward digit span, and the WAB-Repetition subscores). Therefore, an average working memory score for each subject was obtained by averaging standard z-score values for each of the three tests. This average working memory score followed a normal distribution, per Shapiro-Wilk test. There were no significant effects for age, gender, or total brain volume for either of the hypothesized brain regions or for performance on any of the three working memory tasks.

First, our primary hypothesis as to the association between working memory performance and cortical thickness in left pIFS and left pSMG was assessed using one-tailed Pearson bivariate correlation analyses, with a Bonferroni correction applied to account for multiple comparisons resulting in an α level of 0.025. A one-tailed analysis was performed due to the unidirectional hypothesis of reduced working memory performance with cortical thinning in these two ROIs.

Next, an exploratory uncorrected whole-brain analysis was performed to identify additional cortical regions that demonstrate a significant relationship with overall working memory performance. Again, one-tailed Pearson bivariate correlations were performed due to the unidirectional hypothesis of cortical thinning associated with reduced working memory performance. Separate whole-brain one-tailed Spearman correlation analyses were also conducted for each working memory task in order to evaluate task differences. All exploratory correlation analyses used an α level set at 0.05 due to the exploratory nature of the analyses and were performed in IBM SPSS Statistics 25 for Windows.

### Data availability

The analyzed datasets are available from the corresponding author on reasonable request.

## Results

### Brain Atrophy Patterns by Clinical Subtype

Cortical thickness measures in this study revealed differential patterns of left hemisphere atrophy across PPA variants (see Figure 2 for atrophy maps), largely in line with previously described characteristic atrophy in anterior temporal gyri for svPPA patients, in inferior frontal gyrus for nfvPPA patients, and in the temporoparietal junction for lvPPA patients (Rohrer *et al*., 2010; Sapolsky *et al*., 2010; Gorno-Tempini *et al*., 2011; Mesulam *et al*., 2012; Rogalski *et al*., 2014). No significant temporal lobe atrophy was observed in the nfvPPA group.

**Figure 2.**
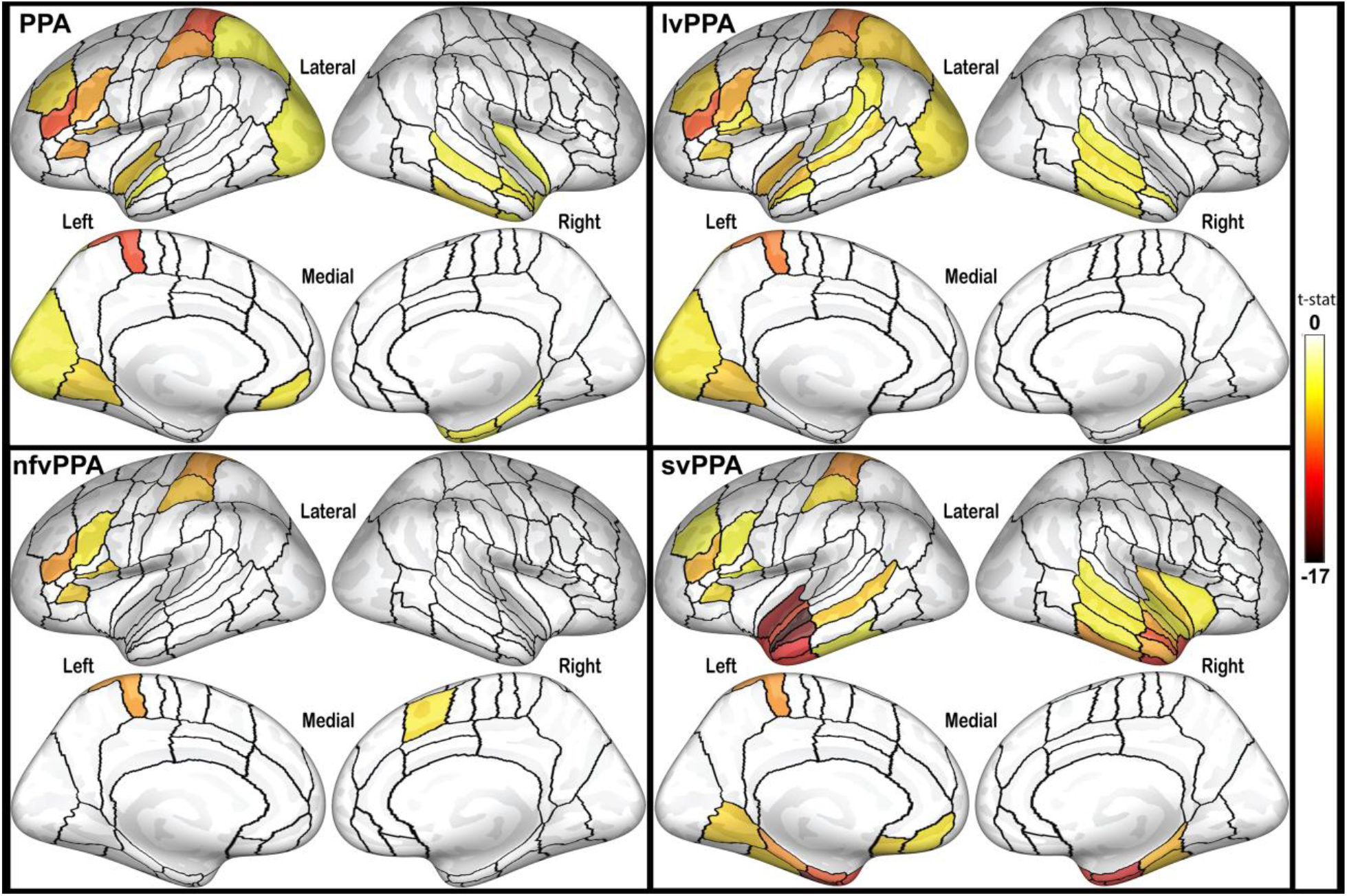
Inflated cortical surfaces showing regions-of-interest with significantly thinner cortex compared to controls. The color scale represents t-statistic of the effect, with false discovery rate correction set at 0.001 for each comparison. PPA = primary progressive aphasia, all variants combined; lvPPA = logopenic variant primary progressive aphasia; nfvPPA = nonfluent variant primary progressive aphasia; svPPA = semantic variant primary progressive aphasia

ANOVA analysis revealed a significant group effect in both left pIFS [F(2, 39) = 5.55, *p* = 0.008] and left pSMG [F(2,39) = 10.46, *p* < 0.001]. Bonferroni post-hoc comparisons revealed a significant difference in left pIFS cortical thickness only between lvPPA and svPPA patients (*p* = 0.006; lvPPA: 2.03 ± 0.15; svPPA: 2.23 ± 0.12; nfvPPA: 2.12 ± 0.21). Differences in left pSMG thickness were present between lvPPA and nfvPPA patients (*p* = 0.002; lvPPA: 2.02 ± 0.05; nfvPPA: 2.30 ± 0.05) and lvPPA and svPPA patients (*p* = 0.001; svPPA: 2.32 ± 0.05).

### Brain-Behavior Correlations

Pearson correlations revealed significant correlations between the average working memory score and cortical thickness in left pIFS (r = 0.397, 95% CI r > 0.155, *p* = 0.005) and left SMG (r = 0.411, 95% CI r > 0.172, *p* = 0.003). Scatter plots displaying cortical thickness in each of these two regions as compared to average working memory scores for each subject are shown in Figure 3.

**Figure 3:**
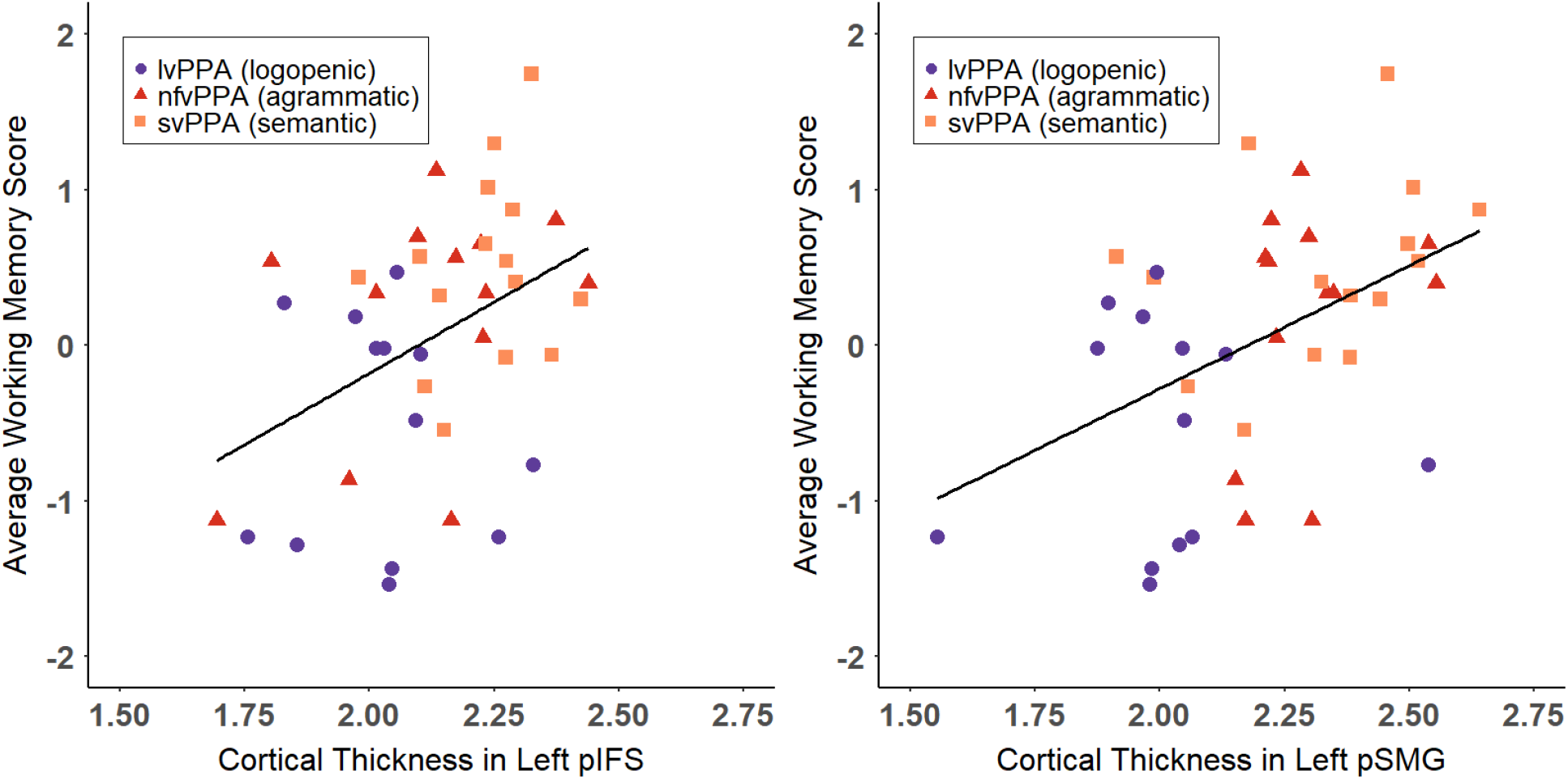
Scatter plot showing relationship between each subject’s mean cortical thickness in left posterior inferior frontal sulcus (pIFS, left panel) and left posterior supramarginal gyrus (pSMG, right panel) as compared to their average working memory score obtained from the mean of z-scores from forward digit span, backward digit span, and the Western Aphasia Battery-Sentence Repetition subscores. Logopenic variant individuals (lvPPA) are shown as purple circles, nonfluent variant individuals (nfvPPA) are shown as red triangles, and semantic variant individuals (svPPA) are shown with orange squares. Solid line shows linear trend for combined group.

Results for exploratory whole brain correlation analyses with the average working memory score, the WAB-Repetition subtest score, forward digit span, and backward digit span are summarized in Table 2, with significant ROIs displayed in Figure 4. Uncorrected results for the average working memory score revealed correlations with cortical thickness in right Heschl’s gyrus (r = 0.415, *p* = 0.003), right posterior dorsal superior temporal sulcus (r = 0.387, *p* = 0.006), left planum temporale (PT; r = 0.385, *p* = 0.006), right anterior middle frontal gyrus (r = 0.412, *p* = 0.007), in addition to the hypothesized regions. WAB-Repetition scores were most strongly correlated with bilateral posterior dorsal superior temporal sulcus (L: r_s_ = 0.461, *p* = 0.001; R: r_s_ = 0.438, *p* = 0.002), left PT (r_s_ = 0.479, *p* = 0.0007), and left posterior superior temporal gyrus (pSTG; r_s_ = 0.452, *p* = 0.001). WAB-Repetition scores were significantly correlated with cortical thickness in left pSMG (r_s_ = 0.341, *p* = 0.014), but not left pIFS (r_s_ = 0.249, *p* = 0.056). Backward digit span was mostly strongly correlated with right Heschl’s gyrus (r_s_ = 0.418, *p* = 0.003) and right frontal regions including superior frontal gyrus (r_s_ = 0.454, *p* = 0.001), anterior dorsal premotor cortex (r_s_ = 0.419, *p* = 0.003), anterior middle frontal gyrus (r_s_ = 0.389, *p* = 0.005), and frontal pole (r_s_ = 0.373, *p* = 0.007). The two hypothesized ROIs were also correlated with backward digit span (pIFS: r_s_ = 0.317, *p* = 0.02; pSMG: r_s_ = 0.282, *p* = 0.035).

**Table 2:**
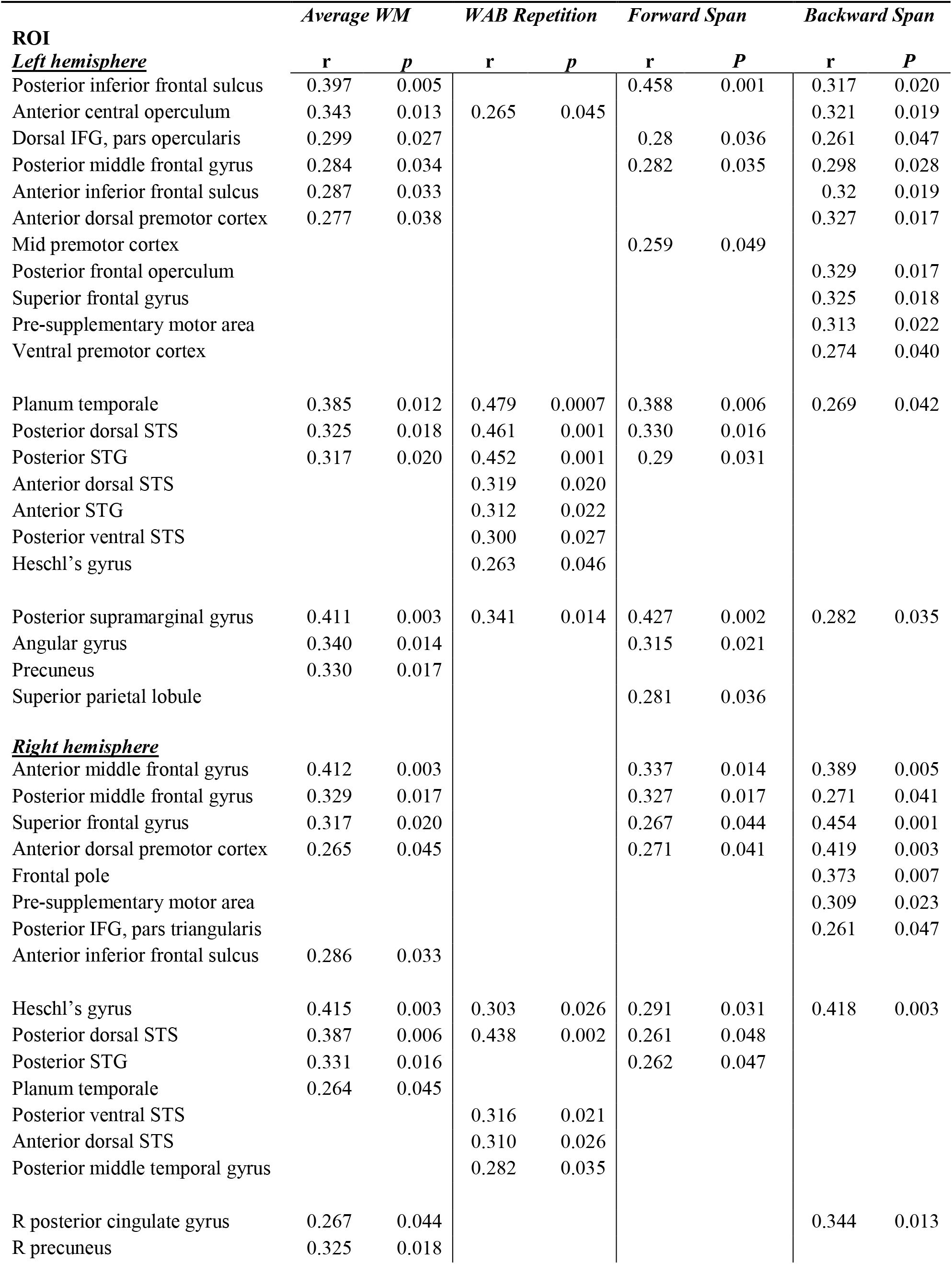
Regions-of-interest (ROIs) with significant (*p* < 0.05) correlation coefficients (r) and *p-values* with each of the three repetition tasks: Western Aphasia Battery (WAB)-Repetition subtest; forward digit span; and backward digit span, as well as ROIs significantly correlated with the average working memory (WM) score, obtained from an average of z-scored values from each of the three individual tests. IFG = inferior frontal gyrus, STG = superior temporal gyrus, STS = superior temporal sulcus

**Figure 4:**
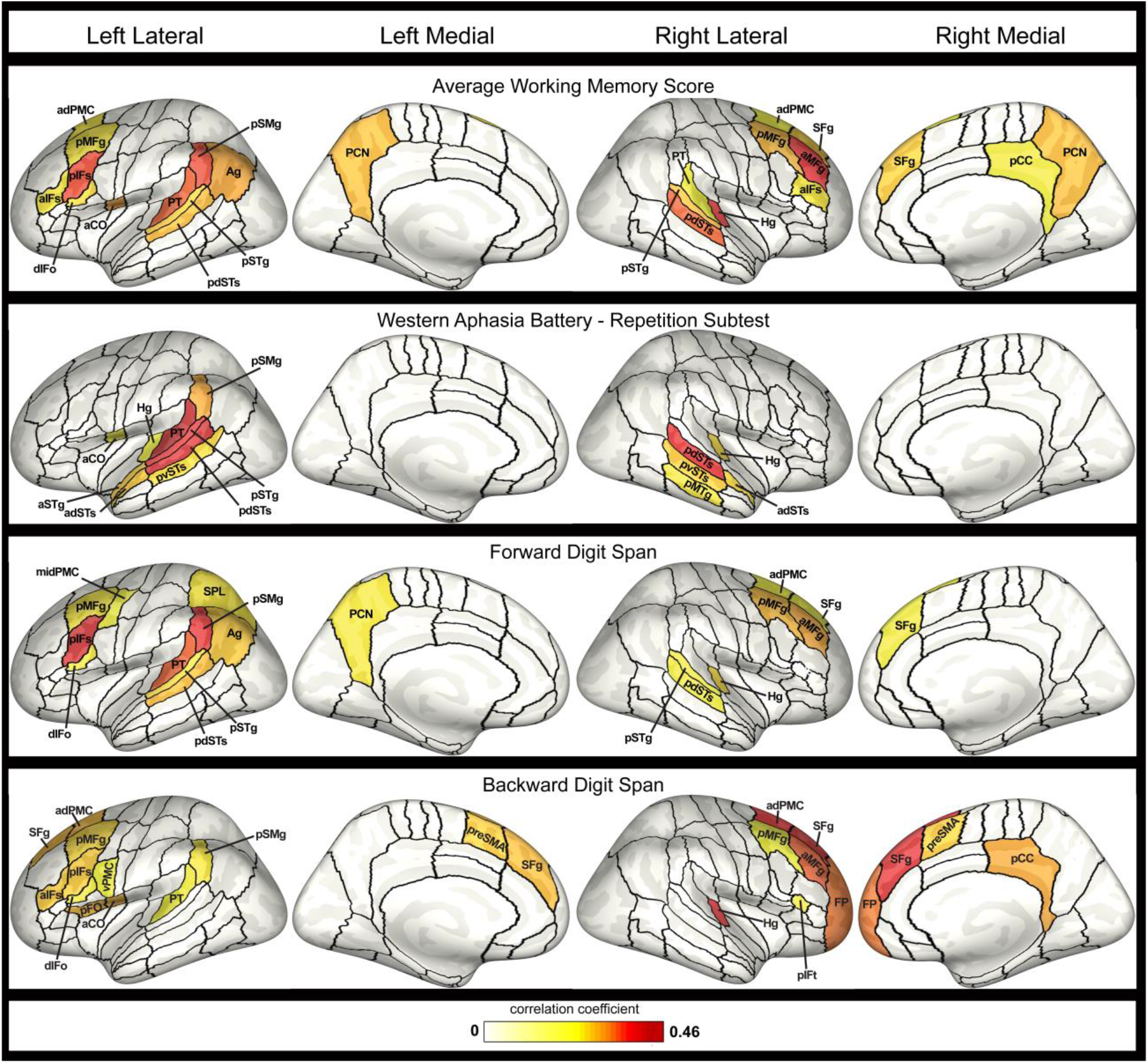
Inflated cortical surfaces showing regions-of-interest (ROIs) with significant correlations with (a) the average working memory score, obtained from an average of z-scored values from each of the three individual tests with each of the three repetition tasks; (b) score on the Western Aphasia Battery-Repetition subtest; (c) forward digit span score; and (d) backward digit span score. Color map reflects the strength of the correlation coefficient, thresholded at *p* < 0.05, uncorrected. Abbreviations: aCO = anterior central operculum, adPMC = anterior dorsal premotor cortex, adSTs = anterior dorsal superior temporal sulcus, Ag = angular gyrus, aIFs = anterior inferior frontal sulcus, aMFg = anterior middle frontal gyrus, aSTg = anterior superior temporal gyrus, dIFo = dorsal inferior frontal gyrus, pars opercularis, FP = frontal pole, Hg = Heschl’s gyrus, midPMC = middle premotor cortex, pCC = posterior cingulate cortex, PCN = precuneus, pdSTs = posterior dorsal superior temporal sulcus, pFO = posterior frontal operculum, pIFs = posterior inferior frontal sulcus, pIFt = posterior inferior frontal gyrus, pars triangularis, pMFg = posterior middle frontal gyrus, pMTg = posterior middle temporal gyrus, preSMA = pre-supplementary motor area, pSMg = posterior supramarginal gyrus, pSTg = posterior superior temporal gyrus, PT = planum temporale, pvSTs = posterior ventral superior temporal sulcus, SFg = superior frontal gyrus, SPL = superior parietal lobule, vPMC = ventral premotor cortex

The strongest correlations with forward digit span were observed in the two hypothesized phonological buffers (left pIFS: r_s_ = 0.458, *p* = 0.001; left pSMG: r_s_ = 0.427, *p* = 0.002), and left PT (r_s_ = 0.388, *p* = 0.006; see Table 2 and Figure 4 for complete results). Because of the large number of ROIs tested in this exploratory analysis, none of these correlations survived FDR-correction for multiple comparisons.

## Discussion

We investigated the neural substrates underlying performance on several clinical tests involving phonological working memory (PWM) by examining the relationship between cortical thickness and behavioral performance in PPA patients. Specifically, we found correlations between average performance across three verbal repetition tasks and cortical thickness in both left pIFS and pSMG in PPA patients. Our results support the involvement of left pIFS in PWM, as proposed by the GODIVA model of speech sequencing (Bohland *et al*., 2010) in which left pIFS serves as an output buffer in addition to a separate phonological buffer in temporoparietal cortex. Exploratory whole-brain analyses revealed distinct brain regions that were correlated with scores from each of the three tasks (in addition to some overlap), demonstrating potential differences in the neural correlates underlying successful performance on sentence repetition, forward digit span, and backward digit span tasks.

### Evidence of a Phonological Content Buffer in Left pIFS

The novel finding of correlations between left pIFS cortical thickness and PWM performance in patients with PPA supports the hypothesized role of left pIFS as a phonological content buffer in the GODIVA model. In this model, left pIFS serves as part of a cortico-basal ganglia-thalamo-cortical planning loop and is specifically responsible for buffering and sequencing the individual phonological units in an upcoming utterance, which are then activated and executed in the correct serial order via projections to ventral premotor cortex for speech output (Bohland et al., 2010; Guenther, 2016). This phonological output buffer should be highly involved in the commonly used tasks analyzed in this study. Specifically, impairment of this phonological output buffer should result in poorer verbal repetition performance due to difficulty buffering and sequencing each phoneme prior to motor execution.

Left inferior frontal regions have long been associated with the articulatory rehearsal component of Baddeley’s working memory model (Paulesu *et al*., 1993; Awh *et al*., 1996; Baddeley, 2003; Baldo and Dronkers, 2006). Within the GODIVA model, left pIFS (in concert with left vPMC) plays a similar role to Baddeley’s articulatory rehearsal process, sequencing through phonological units for either overt speech output or covert rehearsal (Bohland *et al*., 2010). Bohland & Guenther (2006) demonstrated increased activation in left pIFS during production of more complex syllable strings (e.g. increased number of phonemes per sequence), consistent with the region’s proposed function in which additional neural resources are required to code the serial order of additional phonemes. Moreover, a prior meta-analysis of neuroimaging studies identified left pIFS as the only neural region with preferential activation in verbal (compared to nonverbal) working memory tasks (Rottschy *et al*., 2012).

Previous studies of verbal repetition in PPA have identified correlations between repetition deficits and atrophy in temporoparietal regions but not prefrontal regions such as pIFS (Amici *et al*., 2007; Rogalski *et al*., 2011; Leyton *et al*., 2012; Lukic *et al*., 2019). Our study differs from previous work in the use of an ROI-based analysis using the Speech Label parcellation scheme. This parcellation scheme defines subject-specific ROIs to account for inter-subject anatomical variability and allows for finer-scale subdivisions of critical speech cortical regions, like pIFS, improving the localization of speech and language functions for more sensitive statistical analyses (Nieto-Castañon *et al*., 2003; Tourville and Guenther, 2012).

Discrepant findings are also partly explained by differences in the selected repetition tasks; our individual task analyses demonstrated that left pIFS was not significantly correlated with sentence repetition performance on the WAB, consistent with previous literature.

### Evidence of a Phonological Content Buffer in Left pSMG

Average PWM task performance was also significantly correlated with cortical thickness in left pSMG, consistent with prior theoretical accounts of PWM. This finding replicates recent work demonstrating correlations between sentence repetition accuracy and cortical thickness in temporoparietal regions, including left SMG, in patients with PPA (Lukic *et al*., 2019). In lvPPA patients, atrophy in left SMG is correlated with increased phonologic substitution errors (Petroi *et al*., 2020) and with impaired naming, presumably due to phonological impairment (Leyton *et al*., 2012). Similarly, substitution errors and repetition deficits in conduction aphasia have been linked to left SMG damage (Axer *et al*., 2001; Baldo and Dronkers, 2006; Fridriksson *et al*., 2010). Left SMG has also been implicated in functional neuroimaging of phonological working memory tasks as a phonological input buffer or the site of Baddeley’s phonological store (Paulesu *et al*., 1993; Awh *et al*., 1996; Henson *et al*., 2000; Bohland and Guenther, 2006; Rottschy *et al*., 2012; Yue *et al*., 2019). Notably, our exploratory analysis of individual tasks suggests that this buffer extends from left pSMG into the superior temporal lobe, especially PT. This finding is in line with emergent property models where pSTG and PT act as a sensorimotor interface linking acoustic and phonological representations (Jacquemot and Scott, 2006; Hickok and Poeppel, 2007; Buchsbaum and D’Esposito, 2008; Majerus, 2013).

### Task Differences

A distinct set of neural regions (with some overlap) was correlated with each of the three common repetition tasks, with implications for the diagnostic utility of each measure in the clinical management of PPA. This study corroborates previous literature demonstrating the sensitivity of cortical thickness measures to detect subtle deficits in PPA and identify the neural correlates of numerous speech and language domains (Sapolsky *et al*., 2010; Rogalski *et al*., 2011; Cordella *et al*., 2019). Consistent with Rogalski et al. (2011), we found that despite group differences in verbal repetition performance, individual nfvPPA and svPPA patients also presented with subtle repetition impairments (see correlation data in Figure 3), in addition to the more salient deficits in grammar or semantics associated with their primary diagnosis. This study capitalizes on this distribution to analyze the neural substrates of commonly used verbal repetition tasks across PPA variants. Due to the high degree of correlation between the three analyzed tasks, there are a number of regions that were correlated with multiple tasks (e.g., ROIs in the temporoparietal junction) that may reflect a common substrate of PWM necessary across tasks. Not surprisingly, some of these ROIs also overlap with the typical atrophy patterns present in lvPPA, consistent with the hallmark repetition deficits in this population.

Forward digit span performance was most strongly correlated with cortical thickness in the hypothesized phonological content buffers in left pSMG and pIFS. This finding suggests that the forward digit span task may be a purer measure of the function of these phonological buffers, requiring less involvement of higher-level language or cognitive systems than other PWM tasks. Significant correlations between task performance and thickness of adjacent temporoparietal ROIs are consistent with previously reported correlations between left pSTG atrophy and digit span (Leyton *et al*., 2012). Additionally, bilateral middle frontal gyrus (MFG) correlations with both digit span tasks likely reflect this region’s suggested role as part of a multi-domain cognitive system (Niendam *et al*., 2012; Fedorenko *et al*., 2013). Right MFG may also be involved in number manipulation (Menon *et al*., 2000; Zago *et al*., 2008).

The backward digit span task was unique in the contribution of right frontal regions to performance on this task. Bilateral superior frontal gyrus forms part of the frontoparietal control network engaged in sustained attention and executive control (Coull *et al*., 1996; Vincent *et al*., 2008; Niendam *et al*., 2012) with atrophy or lesion shown to impair working memory (Du Boisgueheneuc *et al*., 2006; Barbey *et al*., 2013; Nissim *et al*., 2017). Bilateral preSMA is also considered a core working memory region (e.g., Bohland and Guenther, 2006; Rottschy et al., 2012; Perrachione et al., 2017). Our results support clinical concerns that attentional or executive functioning demands may drive performance on backward digit span tasks more than phonological processing ability (Foxe *et al*., 2013; Beales *et al*., 2019); backward digit span performance was less strongly correlated with cortical thickness in presumed phonology regions, instead requiring intact functioning of more general executive function regions.

Performance on the WAB-Repetition subtest was primarily associated with cortical thickness in left temporoparietal junction. This region is associated with repetition deficits in patients with both post-stroke aphasia and PPA (Damasio and Damasio, 1980; Axer *et al*., 2001; Baldo and Dronkers, 2006; Amici *et al*., 2007; Fridriksson *et al*., 2010; Buchsbaum *et al*., 2011; Rogalski *et al*., 2011; Lukic *et al*., 2019), and it is active during PWM tasks in typical speakers (McGettigan *et al*., 2011; Perrachione *et al*., 2017; Scott and Perrachione, 2019). The identified neural correlates of the WAB-Repetition task also extend into middle and anterior temporal gyri, which may reflect the semantic and syntactic processing involved in the task (Fedorenko *et al*., 2010; Friederici and Gierhan, 2013), with lesions here resulting in comprehension deficits and paragrammatism (Sapolsky *et al*., 2010; Rogalski *et al*., 2011; Turken and Dronkers, 2011; Matchin *et al*., 2020). The involvement of these regions suggests sentence recall may be facilitated by syntactic and semantic knowledge which may outweigh the contribution of the proposed frontal phonological buffer to successful performance on the task (e.g., Baddeley et al., 2009); indeed, the WAB-Repetition task was the only task not significantly correlated with left pIFS. In support of this view, Lukic et al. (2019) found that the use of non-meaningful sentences provided increased diagnostic discrimination as it prevented compensatory use of intact semantic processing that may mask phonologic processing deficits in some lvPPA patients.

### Limitations and Future Directions

In conclusion, the finding of significant correlations between average verbal repetition performance and cortical thickness in both left pIFS and pSMG in a cohort of right-handed PPA patients supports the proposed role of these brain regions. However, we acknowledge several limitations to this work, including the use of only cortical thickness measures in our brain-behavior analyses. White matter and functional connectivity studies have identified disruptions in the structure and function of key speech and language networks in patients with PPA (Galantucci *et al*., 2011; Whitwell *et al*., 2015; Mandelli *et al*., 2016). A more complete picture may emerge by combining multiple structural and functional measures in a single cohort of PPA patients.

Our analyses demonstrate the role of both left pIFS and left pSMG in verbal repetition, but the selected PWM tasks are limited in their ability to sufficiently differentiate deficits in phonological input from output buffer dysfunction, as proposed by prior accounts of PWM. Future work should further isolate and confirm the anatomical correlates of proposed phonological input and output buffers in left pSMG and left pIFS through more precise assessment of deficits in both of these functions. Additionally, our analyses identifying distinct neural correlates of the three repetition tasks are exploratory (i.e., not statistically corrected for the total number of ROIs analyzed), allowing for potential Type I errors. These exploratory results suggest significant task differences that should be further investigated in future work to validate these findings.

## Acknowledgments

We are grateful to our patients and healthy volunteers for their participation in this research.

## Funding

This work was supported by the National Institute on Deafness and other Communication Disorders (NIDCD) of the National Institutes of Health under award numbers R01 DC007683 and R01 DC002852 (PI: F. Guenther), R01 DC014296 (PI: B. Dickerson), and T32 DC013017 (PI: C. Moore).

## Competing Interests

B. Dickerson receives research support from NIH and the Alzheimer’s Drug Discovery Foundation. Dr. Dickerson serves on scientific advisory boards for Acadia, Arkuda, Axovant, Lilly, Biogen, Merck, Novartis, Wave LifeSciences and performs editorial duties with payment for Elsevier (Neuroimage: Clinical and Cortex). Dr. Dickerson also receives royalties from Oxford University Press and Cambridge University Press. The other authors declare no competing financial interests relevant to the manuscript.

## Notes

### Competing Interest Statement

B. Dickerson receives research support from NIH and the Alzheimers Drug Discovery Foundation. Dr. Dickerson serves on scientific advisory boards for Acadia, Arkuda, Axovant, Lilly, Biogen, Merck, Novartis, Wave LifeSciences and performs editorial duties with payment for Elsevier (Neuroimage: Clinical and Cortex). Dr. Dickerson also receives royalties from Oxford University Press and Cambridge University Press. The other authors declare no competing financial interests relevant to the manuscript.

